# AlleleHMM: a data-driven method to identify allele-specific differences in distributed functional genomic marks

**DOI:** 10.1101/389262

**Authors:** Shao-Pei Chou, Charles G. Danko

## Abstract

How DNA sequence variation influences gene expression remains poorly understood. Diploid organisms have two homologous copies of their DNA sequence in the same nucleus, providing a rich source of information about how genetic variation affects a wealth of biochemical processes. However, few computational methods have been developed to discover allele-specific differences in functional genomic data. Existing methods either treat each SNP independently, limiting statistical power, or combine SNPs across gene annotations, preventing the discovery of allele specific differences in unexpected genomic regions. Here we introduce AlleleHMM, a new computational method to identify blocks of neighboring SNPs that share similar allele-specific differences in mark abundance. AlleleHMM uses a hidden Markov model to divide the genome among three hidden states based on allele frequencies in genomic data: a symmetric state (state ‘S’) which shows no difference between alleles, and regions with a higher signal on the maternal (state M) or paternal (state P) allele. AlleleHMM substantially outperformed naive methods using both simulated and real genomic data, particularly when input data had realistic levels of overdispersion. Using PRO-seq data, AlleleHMM identified thousands of allele specific blocks of transcription in both coding and non-coding genomic regions. AlleleHMM is a powerful tool for discovering allele-specific regions in functional genomic datasets.

## Introduction

DNA encodes the blueprints for making an organism, in part by coordinating a complex cell-type and condition-specific gene expression program. Regulatory DNA affects gene expression by controlling the rates of a variety of steps during the transcription cycle, including opening chromatin, decorating core histones and DNA with chemical modifications, initiating RNA polymerase II transcription, and the release of Pol II from a paused state into productive elongation [1]. In addition, mRNAs in most genomes are subjected to a host of post-transcriptional regulatory processes, most of which are influenced by the sequence of the RNA [2]. However, how DNA sequences regulate each step during transcription and the mRNA life-cycle remains poorly understood.

Finding allele-specific differences in the distribution of marks along the genome is one powerful strategy for understanding the link between DNA sequence and the various biochemical processes that regulate gene expression [3,4]. Diploid organisms have two copies of their DNA sequence in the same nuclear environment, providing a rich source of information about how genetic variation affects biochemical processes. Additionally, alleles in a diploid genome share the same environmental signals, cell type-specific differences within a complex tissue, and other potential confounders, making allele-specific signatures a highly rigorous source of information about how DNA sequence affects gene expression.

Despite the general utility allele specific expression, surprisingly few computational methods to detect allelic differences have been proposed. Current methods examine allele-specific enrichment either test SNPs independently [4,5] or combine the location of SNPs using gene annotations [6]. These methods have a number of important limitations. Treating SNPs independently requires a high sequencing depth, and exhibits a bias where regions with higher abundance of the mark of interest are much more likely to be discovered. Summing up the reads within contiguous genomic regions, such as annotated genes, can improve sensitivity and reduce bias by pooling information across SNPs that share the same allelic bias. However, combining reads requires a well-annotated reference genome, which is not available in some species, and also prevents the analysis of marks in unannotated or non-coding regions which are critical for proper genome function.

We developed a hidden Markov model called AlleleHMM to address these limitations. AlleleHMM identifies genomic blocks sharing same allelic bias in a data driven way. AlleleHMM uses the spatial correlation of alleles to identify blocks comprised of multiple SNPs that share the same allele specific differences in mark abundance. We show that AlleleHMM has significantly higher sensitivity and specificity when compared to tests that treat each SNP independently. When applied to publicly available PRO-seq data, AlleleHMM identified thousands of allele specific blocks that lie outside of gene annotations. Thus, AlleleHMM is a powerful new strategy to identify allele-specific differences in functional genomic data.

## Materials and Methods

### Overview of AlleleHMM

The key goal of AlleleHMM is to identify allele-specific blocks of signal in distributed functional genomic data assuming that contiguous genomic regions share correlated allele-specific events. We developed a HMM that represents allelic bias in a distributed genomic mark using three hidden states: symmetric (S) distribution of the mark from both parental alleles (which shows no allelic bias), and maternally- (M) or paternally-biased (P) regions (**Figure 1B**). AlleleHMM takes as input read counts corresponding to each allele, computed using AlleleDB [4,5]. AlleleHMM uses this information to set the parameters of the HMM using Baum Welch expectation maximization. The Viterbi algorithm is then used to identify the most likely hidden states through the data, resulting in a series of candidate blocks of signal with allelic bias. We last calculated the coverage of allele-specific reads count in each predicted AlleleHMM block and performed a binomial test to examine if the block is significantly biased toward maternal or paternal transcription. The last binomial test was performed to eliminate the false positives resulted from multiple counts of a single read that mapped to multiple nearby SNPs.

**Figure 1:**
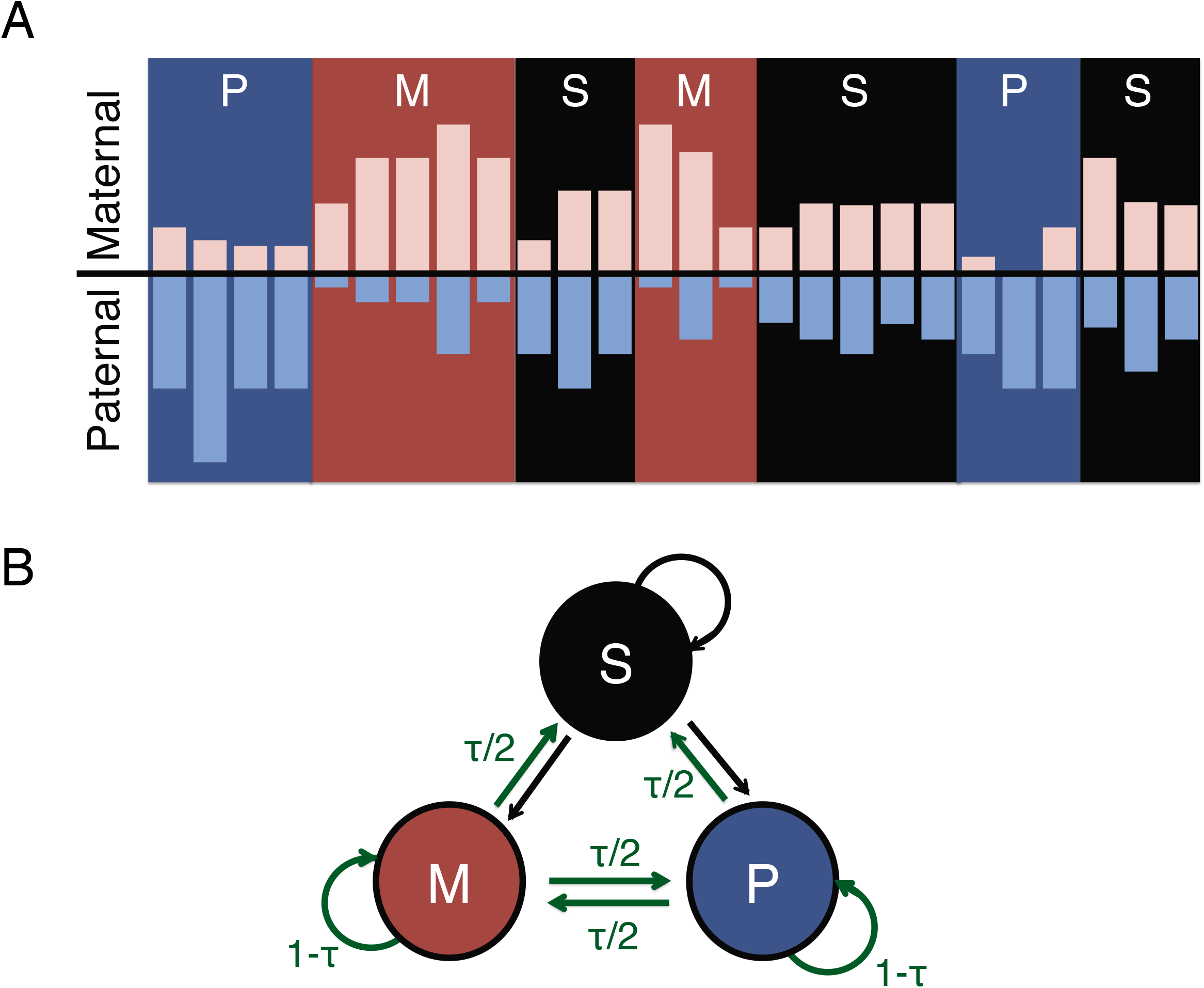
AlleleHMM uses a hidden Markov model (HMM) to infer the allelic bias of genomic markers at each SNPs. (A) Cartoon shows the frequency of reads mapping to the patneral (light blue bars) and maternal (pink bars) allele at positions across the genome (X-axis). Nearby SNPs show similar signatures of allelic bias depicted as blue (P, paternal bias), red (M, maternal bias), or black (S, no evidence of allelic bias) background identified using AlleleHMM. (B) The model structure of AlleleHMM. We model allelic bias using three hidden states: symmetric which shows no allelic bias (S, black), and maternally- (M, red) or paternally-biased (P, blue) regions. SNPs can transition between hidden states. Green arrows represent the transition probabilities set using a user-adjustable tuning parameter, τ.

### HMM structure

There are three hidden states in AlleleHMM (**Fig. 1B**): (S) symmetric transcription which shows no allelic bias, and (M) maternally- or (P) paternally-biased regions. Each state can transit to the other two states or stay in the original state. We used allele-specific read counts of SNPs with at least one mapped read. The distance between SNPs were not considered in the model.

### Transition probability

We used a tuning parameter τ to set the transition probabilities. Without restricting model parameters, the transition probability between states are high and there was a lot of transitions within transcript/gene body. The initial transition probability to other states were set to τ/2, and the initial transition probability to stay in the same state is 1-τ. The transition probability of S states to all states were allowed to change to maximize the likelihood via EM algorithm. The transition probability of M or P states to self and other states are fixed to 1-τ, τ/2, and τ/2.

### Turning parameter optimization using PRO-seq data

We assumed that SNPs within the same transcript are likely to have similar signatures of allele-biased transcription, either symmetric or biased toward one of the parents. Therefore, the optimum value of τ should maximize the fraction of state transitions near transcription start site (TSSs) that are active in that cell type (**Fig. 4A**). We used dREG (discriminative regulatory-element detection from GRO-seq) [7] to predict TSSs from a Precision nuclear run-on sequencing (PRO-seq) dataset from 129/castaneus F1 hybrid mouse embryonic stem cells (mESCs) [8]. Using the same PRO-seq data as input to AlleleHMM, we found that the fraction near TSSs increases monotonically with lower τ and the magnitude of change reduced at τ around 1e-4 to 1e-6 (**Fig. 4B,C**). Therefore, we set τ to 1e-05 to the rest of the study.

### Emission probability

The emission probabilities for all three states were calculated using the binomial distribution given total reads count and maternal reads count on all mapped SNPs ordered by position:

SNP order 1, 2, 3,…,l_c_
Total reads count n_1_,n_2_n_3_,…n_l_.
Maternal reads count x_1_, x_2_, x_3_,…x_l_
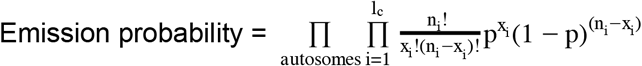

### Learning procedure for transition and emission probabilities

We use the Baum–Welch algorithm [9,10], an expectation maximization (EM) algorithm that utilizes forward-backward algorithm [10] to learn the transition and emission probability in the AlleleHMM model. Then we utilized Viterbi decoding [10] to learn the most likely hidden state sequence.

### Identify allele-specific transcribed blocks

Nearby SNPs sharing the same hidden state were stitched into blocks and then we calculated the coverage of reads in each block as follows: Reads were mapped to diploid genome using bowtie as coded in AlleleDB pipeline [5]. The bowtie output, including reads and their mapping position, were further separated to maternal- and paternal-specific file. Then, the coordinates were transferred to the appropriate reference genome (mm10 or hg19) using liftOver. We used bedtools coverage to calculate the number of reads falling into each stitched block. Binomial tests were performed for each blocks and false discovery rate was estimated to account for multiple testing. Thess steps were performed to eliminate the false positives from multiple counts of a single read that mapped to multiple nearby SNPs. It was also used to evaluate if the tuning parameter τ was chosen appropriately. When τ was chosen appropriately, the percentage of the blocks removed was minute, 0.43% for PRO-seq data from the 129/castaneus F1 hybrid mESCs and 1.10% for the GRO-seq of GM12878. AlleleHMM outputs two bed files: one with all blocks and the other only reports the blocks with a FDR <= 10% as significantly allelic biased.

### Performance test with synthetic data

To test how AlleleHMM performs compared with current standards in the field, which involves testing SNPs independently, we developed a simulation strategy where the location of allele-biased blocks is known. The synthetic data was composed of three blocks, each representing a region with allelic bias as shown in on the top of Figure 2. While the flanking blocks remain allelic balanced and consistent, the following parameters were changed to simulate the middle block with allele-biased transcription: length, expression level, and the degree of allele imbalance. Length was defined as the number of continuous SNPs sharing same allele specificity, and was set to 100 when testing other parameters. We chose the parameters of our simulation based on the length of a typical mammalian gene and a heterozygosity similar to 129/castaneus F1 hybrid mouse, which had approximately 140 SNPs per gene including introns. Expression level, or the average read count per SNP in the block, was set to 10 when testing other parameters. The degree of allele imbalance was the probability that a read is coming from the maternal allele (matP) in a binomial event or beta-binomial event, and was set to 0.9 when testing other parameters. The total read counts of each location was simulated with poisson distribution and the allele-specific read counts was simulated by binomial distribution or beta-binomial distribution with overdispersion of 0.25. The overdispersion of 0.25 was chosen based on the estimates of two real data sets: GRO-seq of GM12878 and PRO-seq of 129/castaneus F1 hybrid mESCs. The estimate was performed in R using VGAM library with all SNPs covered by at least 5 reads.

**Figure 2:**
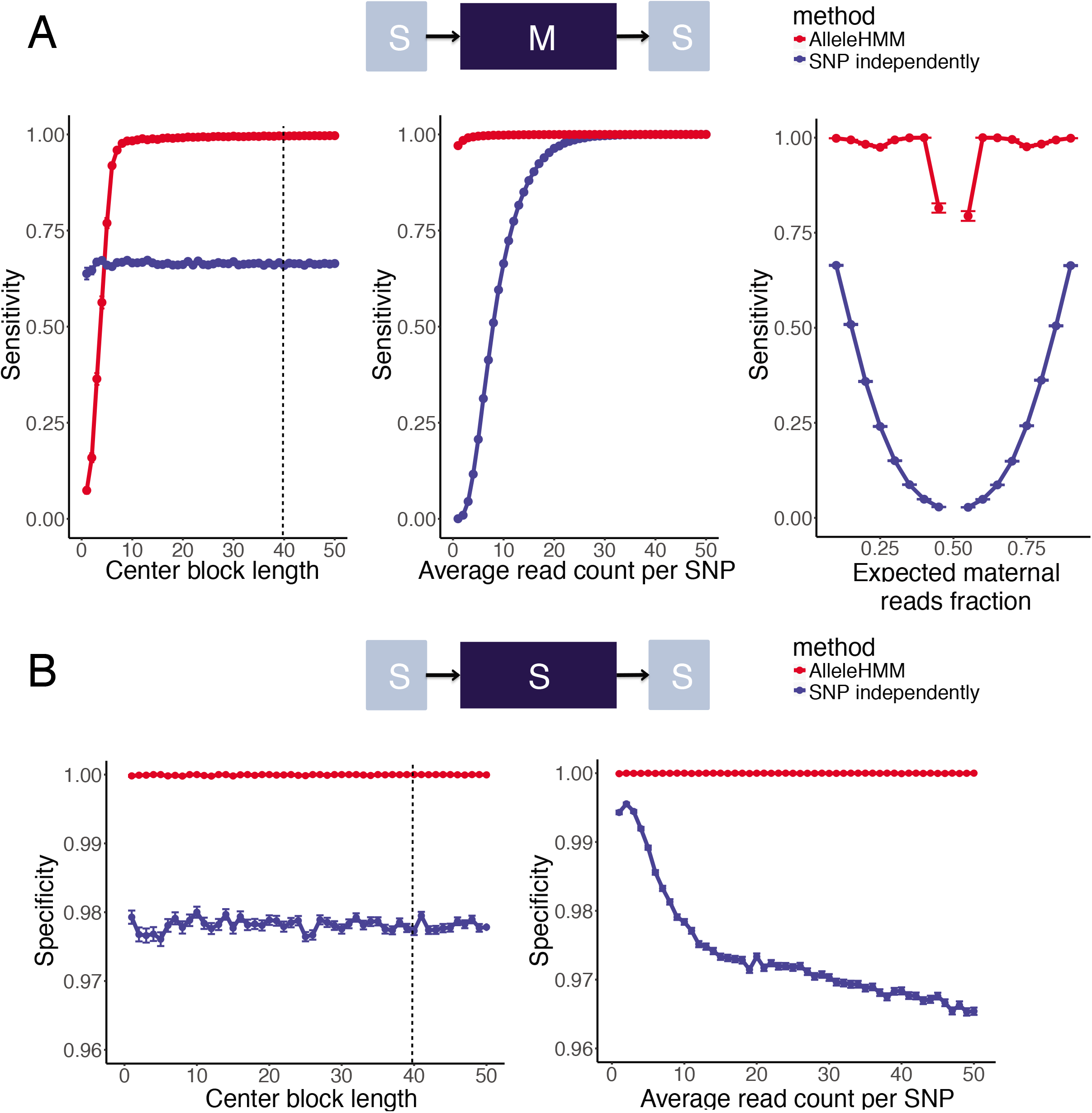
AlleleHMM had better sensitivity and specificity compared with the naive standard practice of performing a binomial test for each SNP independently. (A) Scatterplots show the sensitivity for each SNP in the center block of AlleleHMM (red) and independent binomial tests (blue) as a function of the length of a maternally-biased block (left), the average read count at each SNP (center), or the expected maternal fraction (right). Error bars represent the standard error of 1000 independent simulations. The dotted line indicates the average number of SNPs per human gene. (B) Scatterplots show the specificity of AlleleHMM (red) and independent binomial tests (blue) as a function of the length of a maternally-biased block (left) or the average read count at each SNP (right). Error bars represent the standard error of 1000 independent simulations. Dotted line indicates an estimate number of SNPs per human gene.

### Performance test with real biological data

We applied AlleleHMM and an independent binomial test implemented in AlleleDB on GRO-seq of GM12878 and PRO-seq of 129/castaneus F1 hybrid mESCs. AlleleHMM blocks were compared to the GENCODE gene annotations. We used Release 28 (mapped to GRCh37) for GM12878 and Release M17 (GRCm38.p6) for 129/castaneus F1 hybrid mouse. The correspondence between AlleleHMM blocks and GENCODE gene annotations were performed by bedtools and then summarized using in-house script.

To test the correlation between transcription and H3K27me3 in GM12878, we mapped the H3K27me3 ChIP-seq reads to the diploid genome of GM12878 using bowtie as coded in AlleleDB [5]. The bowtie output, including reads and their mapping position, were further separated to maternal- and paternal-specific file. Then, the coordinates were transferred to the reference genome (hg19) using liftOver. We used bedtools coverage to calculate the number of H3K27me3 ChIP-seq reads falling into each AlleleHMM block obtained from GRO-seq of GM12878. We then calculated the ratio of maternal-specific and paternal-specific H3K27me3 ChIP-seq reads in each GRO-seq AlleleHMM block and summarized using in-house R scripts.

### Data used in this study

PRO-seq of 129/castaneus F1 hybrid mouse embryonic stem cells: SRA ID number SRR4041366.

GRO-seq of GM12878: SRA ID number SRR1552485

H3K27me3 ChIP-seq data of GM12878 were fastq files from ENCODE: ENCFF000ASV, ENCFF000ASW, ENCFF000ASZ, ENCFF001EXM, ENCFF001EXO

## Results

### Finding allele specific differences using a hidden Markov model

Biochemical marks indicative of genome function are frequently spread across broad genomic intervals. Differences in mark abundance between two heterozygous alleles are therefore often correlated across multiple adjacent single-nucleotide polymorphisms (SNP) (**Fig. 1A**). We developed AlleleHMM to identify genomic regions that share allele-specific differences in functional mark abundance. AlleleHMM takes as input counts of reads mapping unambiguously to each of the two alleles in heterozygous positions of a phased reference genome. AlleleHMM models the data using a hidden Markov model (HMM) that divides the genome among three hidden states: a symmetric state (state ‘S’) which shows no allelic difference in mark abundance, and regions with a higher signal on the maternal (M) or paternal (P) allele (**Fig. 1B; see methods**). AlleleHMM models the distribution of read counts mapping to each allele using a binomial distribution. To limit switches between states to those which are best supported by the underlying data, we introduced a user-adjustable tuning parameter, τ, that constrains the transition probability out of either the maternal or paternal state. Aside from τ, all other model parameters are set using expectation maximization over the provided data.

### Performance test with binomial-distributed simulated data

To determine how AlleleHMM performed in practice, we simulated blocks of contiguous SNPs where the allelic imbalance was known. We simulated a sequence of SNPs composed of three blocks using the binomial distribution (**Fig. 2A, top**): two blocks with equal signal in both alleles and one middle block that exhibited a known difference in signal between alleles. Although AlleleHMM is applicable to any type of functional genomic sequencing data, we chose simulation parameters characteristic of precision nuclear run-on and sequencing (PRO-seq) [11], a direct measurement of RNA polymerase. PRO-seq data has a low background, but typically only a few reads map to each SNP, resulting in poor power to identify differences. Moreover, since RNA polymerase densities on each allele are determined by events occurring in the gene promoter [1,12], allele specific differences should generally be shared between all SNPs within the same transcription unit.

We evaluated the performance of AlleleHMM after we systematically changed the length, signal level, and degree of allelic imbalance in the middle block holding other parameters constant (see methods) (**Fig. 2A**). AlleleHMM identified allelic differences in signal intensity of simulated data with higher sensitivity and specificity compared with simple methods that perform independent binomial tests at each SNP (**Fig. 2**). The sensitivity of AlleleHMM for each simulated allele-specific SNP in the center block increased with block length. AlleleHMM had a higher sensitivity than independent binomial tests when the center block contained as few as 5 adjacent SNPs, much shorter than observed in most mammalian transcription units (on average, 39.7 SNPs per gene for human CEPH Utah and 237 SNPs for 129/castaneus F1 hybrid mouse [8], **Fig. 2A, left**). AlleleHMM had a higher sensitivity across the spectrum of signal levels (**Fig. 2A, center**), and was much more sensitive to allele-specific differences in signal intensity that were smaller in magnitude between alleles (AlleleHMM sensitivity nearly 1.0 for bias <0.4 or >0.6 compared to <0.1 for independent SNPs **Fig. 2A, right**). Notably, AlleleHMM had a higher specificity throughout the range of expression and block length parameters than treating SNPs independently (**Fig. 2B**), demonstrating that AlleleHMM does not trade a higher sensitivity for a higher false discovery rate. Simulation tests using parameters fixed to the averages in a typical human genome revealed that AlleleHMM also outperformed independent binomial tests in genomes with a lower heterozygosity (**Supplementary Fig. 1**). Thus we conclude that AlleleHMM had better sensitivity and specificity for allele-specific-transcription in synthetic data simulated using the binomial distribution.

### Performance test with overdispersed synthetic data

Many short-read datasets exhibit overdispersion due to a variety of technical factors, which increases false discovery rates for identifying allele-specific differences [5]. To test how AlleleHMM performed with overdispersed data, we applied a similar simulation strategy using a beta-binomial distribution to simulate read counts with varying degrees of overdispersion. AlleleHMM had a reasonably high sensitivity and specificity across the spectrum of distinct overdispersion values (**Fig. 3A**). AlleleHMM retained a sensitivity >0.95 and specificity near 1.0 at realistic overdispersion levels estimated using two independent GRO-seq/PRO-seq datasets: human GM12878 lymphoblastoid cells (overdispersion of 0.24) [13] and 129/castaneus F1 hybrid mouse embryonic stem cells (mESCs) (overdispersion of 0.26) [8]. AlleleHMM outperformed the independent application of binomial tests throughout the range of overdispersion levels (**Fig. 3A**).

**Figure 3:**
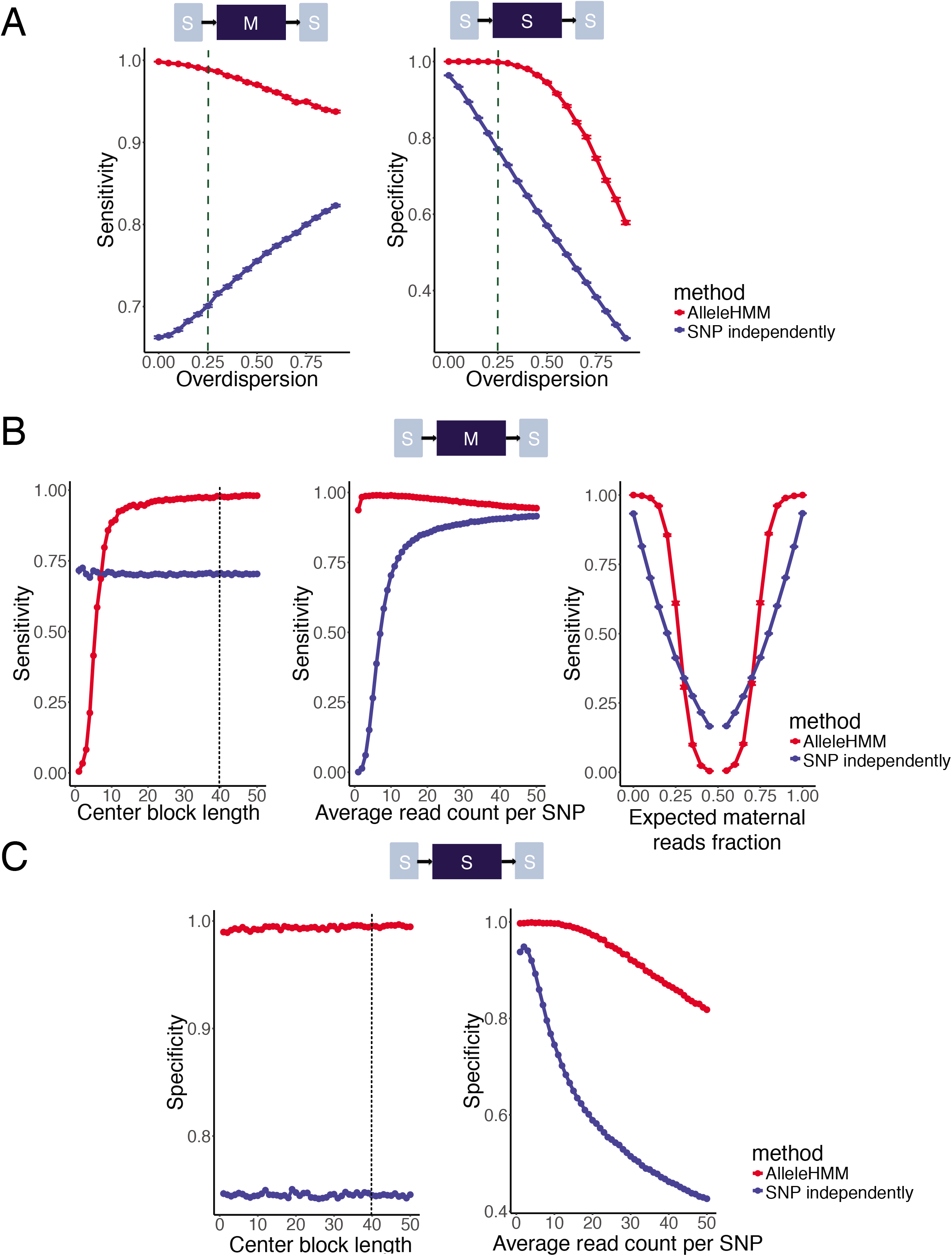
AlleleHMM reduced false discovery rates in overdispersed data. (A) Scatterplots show sensitivity (left) and specificity (right) of AlleleHMM (red) and independent binomial tests (blue) as a function of the overdispersion parameter in beta-binomial distributed simulated data. Error bars represent the standard error of 1000 independent simulations. Dashed lines indicate the mean of overdispersion estimated from GRO-seq of GM12878 and PRO-seq of 129/castaneus F1 hybrid mESCs. (B) Scatterplots show the sensitivity of AlleleHMM (red) and independent binomial tests (blue) as a function of the length of a maternally-biased block (left), the average read count at each SNP (center), or the expected maternal fraction (right) with an overdispersion of 0.25. Error bars represent the standard error of 1000 independent simulations. The dotted line indicates an estimate number of SNPs per human gene. (C) Scatterplots show the specificity of AlleleHMM (red) and independent binomial tests (blue) as a function of the length of a maternally-biased block (left) or the average read count at each SNP (right) with an overdispersion of 0.25. Error bars represent the standard error of 1000 independent simulations. The dotted line indicates an estimate number of SNPs per human gene.

To test how AlleleHMM performance varied with the length, signal level, and the degree of allelic imbalance when the input data was overdispersed, we fixed overdispersion to 0.25 (dashed lines in **Fig. 3A**) and performed simulation experiments similar to those described for the binomial distribution, above. AlleleHMM sensitivity increased with block length, and was higher than independent binomial tests with as few as 8 adjacent SNPs (**Fig. 3B, left**). AlleleHMM was also highly sensitive to a range of allelic bias <0.2 or >0.8 (**Fig. 3B, right**). AlleleHMM had a higher specificity than independent binomial tests across the spectrum of length (the number of SNPs per gene, **Fig. 3C, left**) and signal levels (average read counts per SNP, **Fig. 3C, right**). The specificity of both AlleleHMM and independent binomial tests declined as read count increased, but the rate of decrease was lower for AlleleHMM (**Fig. 3C, right**). Moreover, AlleleHMM exhibited a high sensitivity within the range at which it maintained a high specificity (2-20 reads supporting each SNP, Fig. 3B, center), suggesting that subsampling highly expressed regions may be a viable strategy to deal with overdispersion in practice. Taken together, AlleleHMM reduced the false discovery rate that comes with overdispersion while it maintained a higher sensitivity at realistic parameters taken from PRO-seq datasets.

### AlleleHMM identifies widespread allele specific transcription using PRO-seq data

We applied AlleleHMM to two public run-on and sequencing datasets: PRO-seq data from 129/casŕaneus F1 hybrid mESCs and GRO-seq from a human GM12878 lymphoblastoid cell line. AlleleHMM defined a user-adjustable tuning parameter, τ, that limits the frequency of switches between states. To determine the optimum value of τ for PRO-seq data, we assumed that switches in allele specificity should generally arise near a transcription start site (TSSs) (**Fig. 4A**). We evaluated the proportion of AlleleHMM blocks that start within a fixed distance of a TSS defined using dREG [7,14] over a range of τ in the mESC dataset. As expected, as τ increased, a larger fraction of AlleleHMM blocks occur within a defined distance of a dREG annotated TSS (**Fig. 4B**). As τ approached ~1e-05, the fraction of AlleleHMM blocks beginning within 5kb of a dREG site saturated at ~50% (**Fig. 4B black line, Fig. 4C**). In analyses that follow we therefore fixed τ to 1e-05.

**Figure 4:**
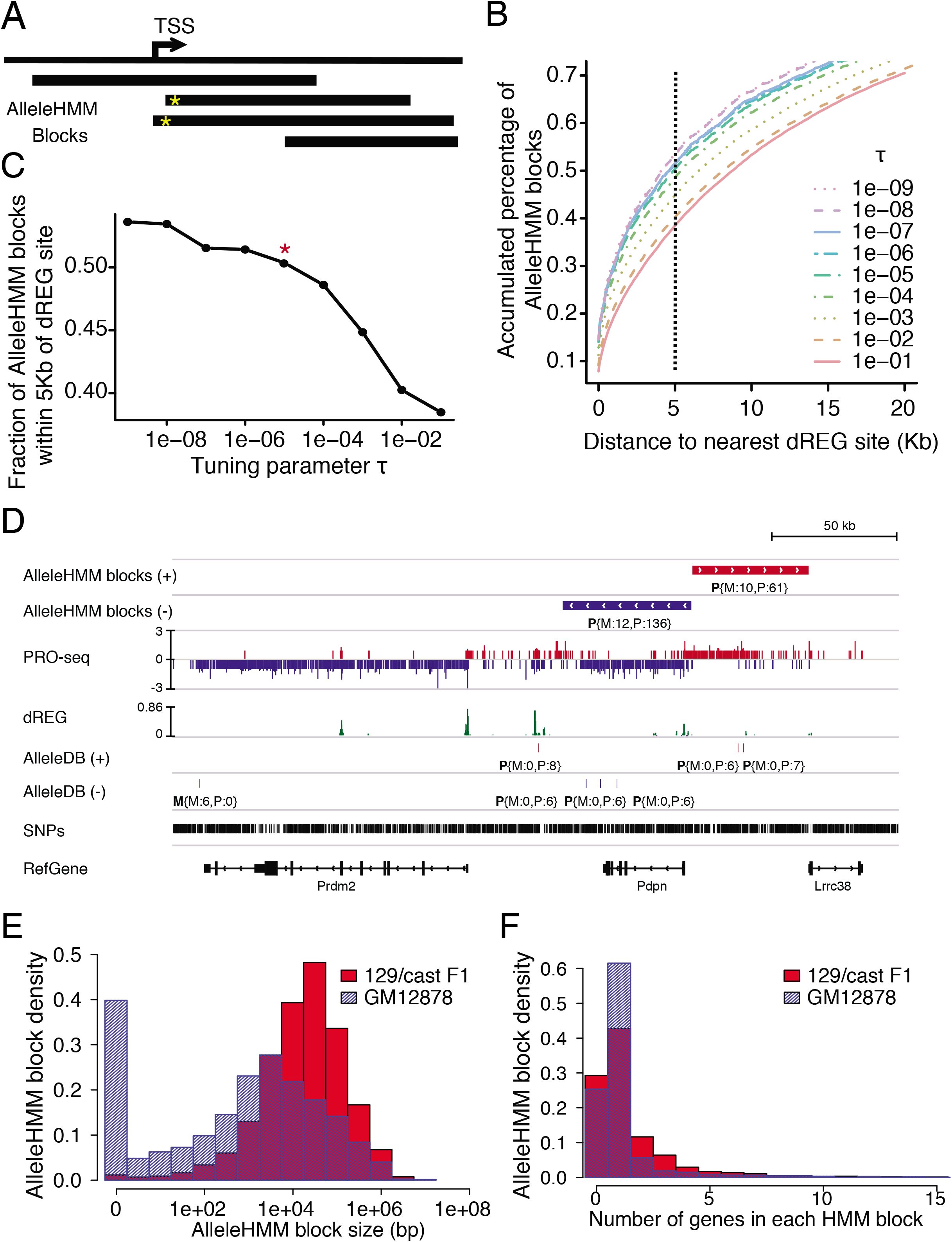
Application of AlleleHMM to PRO-seq and GRO-seq data. (A) Cartoon illustrating how we set the tuning parameter τ. We assumed that SNPs within the same transcript have similar signatures of allelic bias. Therefore, the optimum value of τ should maximize the fraction of state transitions near a transcription start site (TSSs) that are active in that cell type. The black bars are AlleleHMM blocks, represents a region with significant allelic bias. Those with yellow star have state transitions near TSSs. (B) Plot shows the distance between the beginning of AlleleHMM blocks and its closest TSS identified using dREG for PRO-seq data from a 129/castaneus F1 hybrid mouse. Different lines indicate AlleleHMM blocks predicted using different values of the tuning parameter, τ. (C) Scatterplot shows the fraction of AlleleHMM blocks within 5Kb of the nearest TSS predicted by dREG (Y-axis) as a function of the tuning parameter τ (X-axis). The red star indicates a value near a point of saturation (τ = 1e-5) used for the remainder of this study. (D) Genome browser view shows the application of AlleleHMM and independent binomial tests (implemented in AlleleDB) to PRO-seq data from a 129/castaneus F1 hybrid mouse. The allele-specific reads count of the blocks and SNPs are denoted as P{M:12,P:136}, meaning that the block is paternal-biased (P) with 12 maternal-specific (M) reads and 136 paternal-specific (P) reads. (E) Histograms show the distribution of AlleleHMM block size of PRO-seq data from a 129/castaneus F1 hybrid mouse (red) and GRO-seq data from a human cell line GM12878 (blue) in log scale (X-axis). (F) Histograms show the fraction of AlleleHMM blocks as a function the number of genes it contains. PRO-seq data from a 129/castaneus F1 hybrid mouse is in red and GRO-seq data from a human cell line GM12878 is in blue.

Running AlleleHMM genome-wide revealed thousands of regions with maternal or paternal-specific RNA polymerase abundance. AlleleHMM identified 3,483 and 4,026 ‘blocks’ with significant allele-specific differences in mESCs and GM12878, respectively. As illustrated by allele-specific blocks of signal overlapping *Pdpn* and its upstream antisense transcription unit (**Fig. 4D**), AlleleHMM often identified blocks that started near TSSs, and extend across multiple SNPs that in many cases spanned a gene annotation (**blue AlleleHMM block in Fig. 4D**). The average genome size of each block in the F1 hybrid dataset was ~166 kb (**Fig. 4E**). Blocks were larger in the mESC dataset than in GM12878, owing to a combination of differences in heterozygosity and sequencing depth between datasets (**Supplementary Fig. 2**). Approximately 25% of AlleleHMM blocks did not contain any GENCODE gene annotation (**Fig. 4F; example in Fig. 4D**), for example the antisense transcription unit upstream of *Pdpn* (**red AlleleHMM block in Fig. 4D**). Many AlleleHMM blocks contained more than one gene annotation (28% of F1 hybrid mouse blocks and 13% of GM12878 blocks), indicating groups of nearby genes that share similar allelic differences in expression.

### Comparison between AlleleHMM and standard statistical methods

We compared blocks identified using AlleleHMM in PRO-seq data to SNPs identified using independent binomial tests. Surprisingly, few SNPs were identified as allele specific using both AlleleHMM and independent binomial tests on each SNP. In F1 hybrid mESCs, for example, AlleleHMM identified 153,543 heterozygous SNPs with 1 or more read in 3,483 AlleleHMM blocks. Only 9,636 of the SNPs identified using AlleleHMM were also discovered using independent binomial tests (~6% of SNPs; **Fig. 5A, top**). Likewise, in GM12878, AlleleHMM identified 32,599 SNPs in 4,026 blocks, while only 13,086 SNPs were identified using both methods (**Fig. 5A, bottom**). As expected, SNPs identified by AlleleHMM in PRO-seq data largely reflect heterozygous positions covered by too few reads to confidently assign allele specificity when treating SNPs independently, whereas those identified using both methods had a higher read depth (**Supplementary Fig. 3**). Nevertheless, blocks of SNPs identified by AlleleHMM make biological sense when considered across an entire transcription unit (**Fig. 4D**). Taken together, these observations are consistent with AlleleHMM making substantial improvements in sensitivity for allele specific differences in genes with lower expression levels.

**Figure 5:**
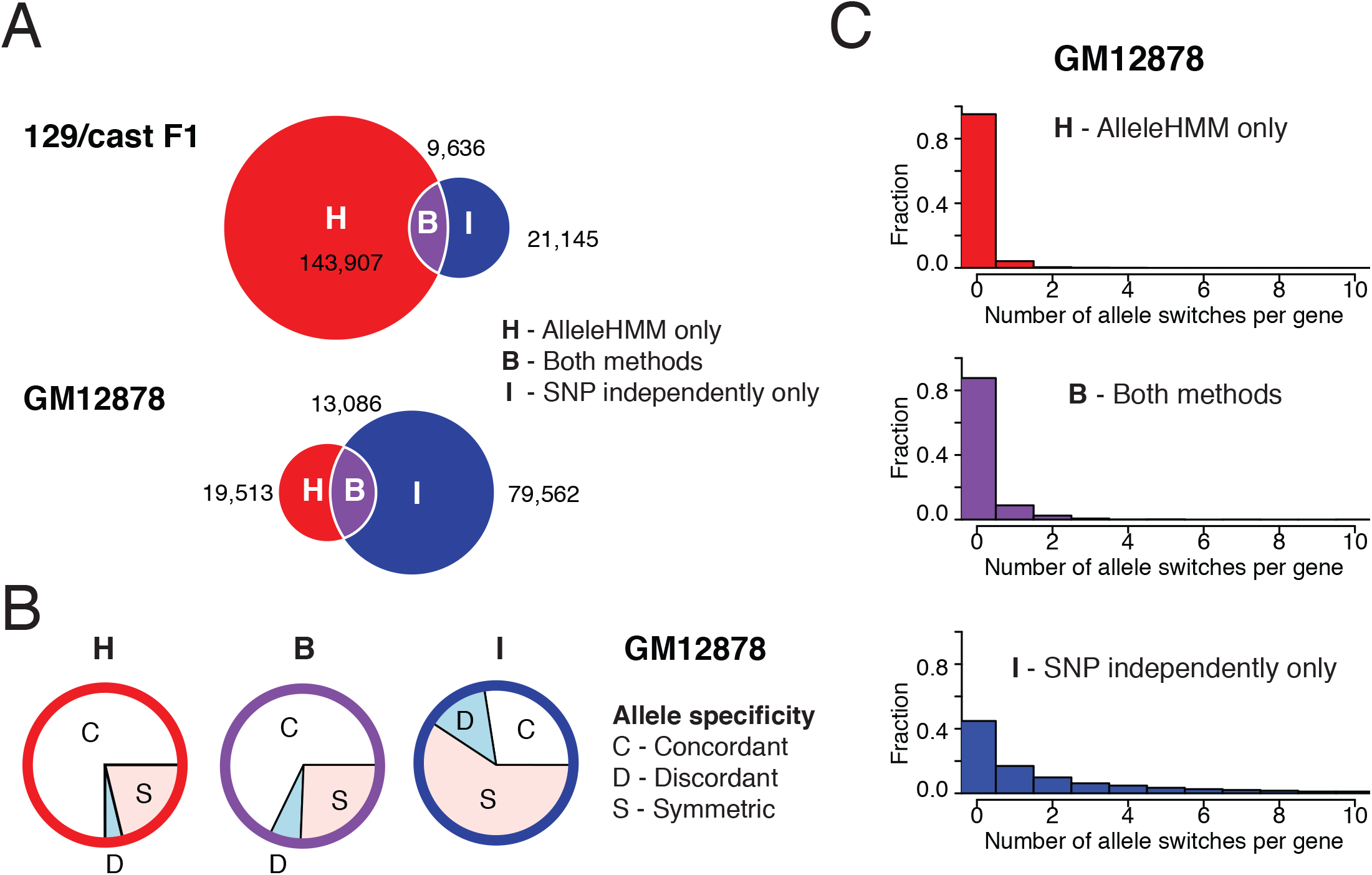
Comparison between AlleleHMM and independent binomial tests. (A) Venn diagrams show the number of allele-specific SNPs identified by AlleleHMM only (H, red), independent binomial tests (I, blue, implemented in AlleleDB), and intersection of both methods (B, purple) from PRO-seq data of a 129/castaneus F1 hybrid mouse and GRO-seq data from a human cell line GM12878. (B) Pie charts show the proportion of allele-specific SNPs in GM12878 GRO-seq data that are within genes having no evidence of allele specificity over the gene (symmetric, S, pink), or genes that show higher expression on the same (concordant, C, white) or the opposite (discordant, D, light blue) haplotype. (C) Histograms show the fraction of genes as a function the number of allele-specificity switches the gene contains. Allele-specificity was determined by AlleleHMM only (H, red, top), independent binomial tests (I, blue, bottom), and intersection of both methods (B, purple, middle) using GRO-seq data from a human cell line GM12878.

We were also surprised to find large numbers of SNPs reported as allele-specific using independent binomial tests without a corresponding discovery by AlleleHMM (n= 21,145 [mESC] or 79,562 [GM12878]). To investigate whether these SNPs were false negative calls by AlleleHMM or false positives by the binomial test, we asked whether SNPs identified as having significant allele specificity within a single gene annotation generally shared the same allele specificity as the gene annotation overall. Because RNA polymerase in the gene body is controlled in the promoter region, we expected bona-fide cases where a single SNP disagrees with the annotation to be rare. Consistent with our expectation, the majority of allele-specific SNPs identified by AlleleHMM were concordant with the direction of changes in allele specificity when SNPs were combined across the entire gene annotation (**Fig. 5B, concordant [C] in white**). By contrast, SNPs identified as allele-specific using independent binomial tests were most often identified within gene annotations that were not allele specific when SNPs were combined across the entire gene (**Fig. 5B, symmetric [S] in pink**). Moreover, highly expressed genes frequently had multiple SNPs within the gene called as allele-specific using independent binomial tests, but often the direction of allele specificity switched across the annotation (**Fig. 5C, bottom**), resulting in no evidence of allelic bias when SNPs within annotations were merged. By contrast, AlleleHMM identified a single block covering annotations in >80% of cases (**Fig. 5C**). Taken together, these results suggest that many of the SNPs identified only by independent binomial tests are likely to be false positives.

### Allele-specific transcription negatively correlates with allele-specific silencing marks

To find further independent validation for blocks of allele-specific transcription, we asked whether we could recover the negative relationship expected between transcription and histone marks associated with transcriptional repression, especially H3K27me3. We focused on GM12878, for which there is publicly available ChIP-seq data profiling the distribution of H3K27me3 [15,16]. Mapping H3K27me3 ChIP-seq data onto AlleleHMM blocks identified using GRO-seq revealed 113 with a significant allelic imbalance in H3K27me3 ChIP-seq data. As expected, the degree of allelic imbalance in H3K27me3 ChIP-seq was inversely correlated with that of GRO-seq (Pearson’s R =-0.57; **Fig. 6**). The slope of the best fit line implies that a 2-fold change in H3K27me3 was associated with ~5.7-fold change in transcription. Assuming a similar dynamic range in both assays, this result implies that relatively modest changes in H3K27me3 may have a relatively large average impact on transcription. Thus, AlleleHMM reveals blocks of allelic bias which are largely in agreement with orthogonal genomic assays.

**Figure 6:**
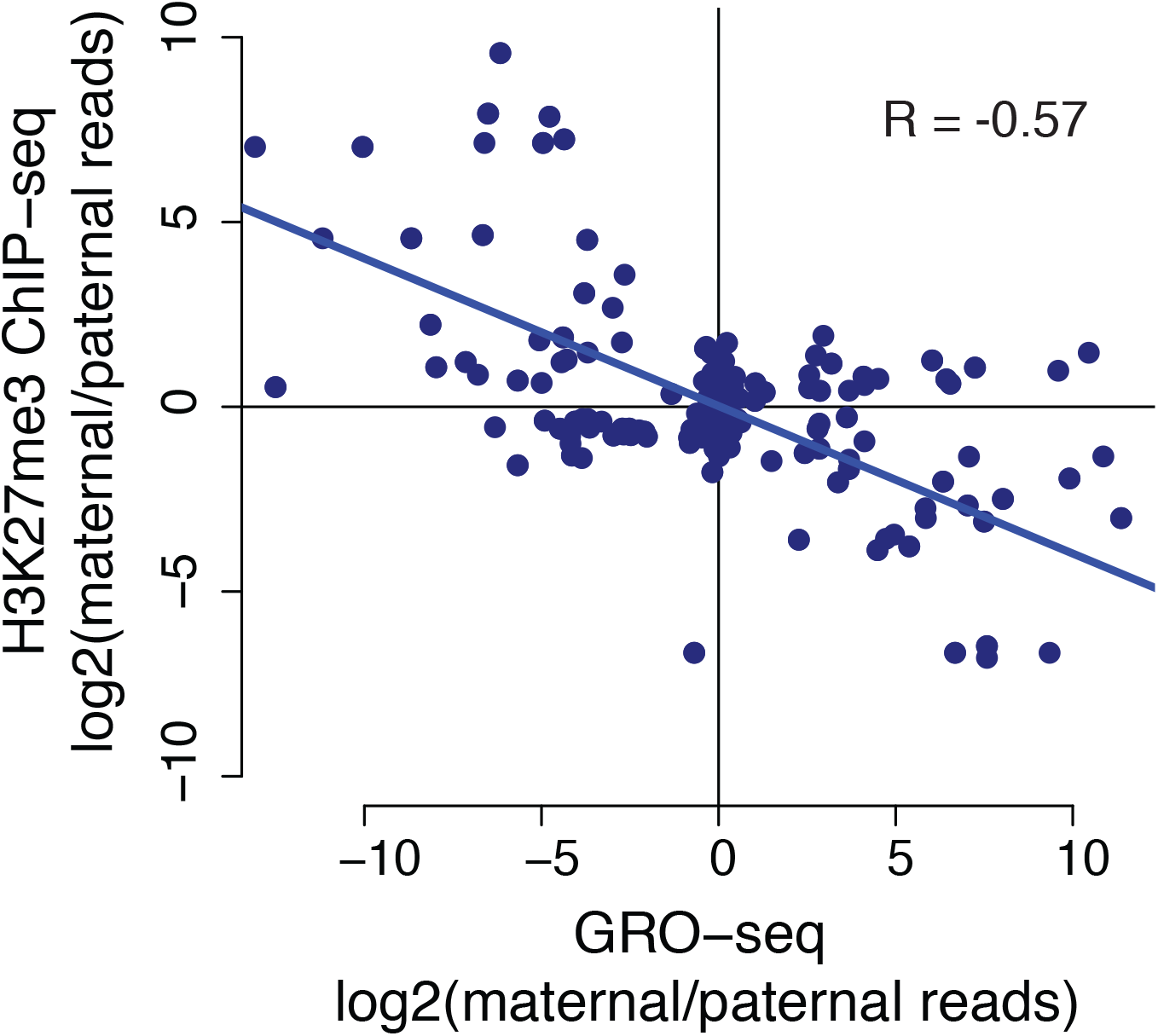
Allele specific transcription correlates with allele specific H3K27me3. Scatterplots show the allelic imbalance of H3K27me3 ChIP-seq data (Y-axis) as a function of allelic imbalance in GRO-seq data (X-axis). The trendline shows the best fit based on a total least squares regression. The Pearson correlation is shown on the plot (R = -0.57, p-value < 2.2e-16). The slope of the best fit line is −2.5.

## Discussion

Here we have introduced AlleleHMM, a new HMM-based tool to identify allele specific differences in functional genomic marks. AlleleHMM uses the spatial correlation in allele-specific differences in a phased diploid genome, providing substantially higher sensitivity and specificity for differences than alternative approaches that use independent statistical tests for each SNP. Likewise, AlleleHMM allows the detection of allele specific differences in transcription that lie outside of annotations, or that are not consistent with annotated genes, allowing AlleleHMM to measure allelic bias in non-coding regions or in poorly annotated genomes.

We demonstrated both a higher sensitivity and specificity under a range of assumptions using simulation tests that are indicative of real sequencing data. Furthermore, analysis of PRO-seq (mESCs) and GRO-seq (GM12878) data suggested that our approach has a higher sensitivity and specificity than alternative approaches. We demonstrated major benefits in sensitivity for more weakly expressed genes and for more subtle differences in mark abundance between alleles, resulting in the discovery of sites that are substantially less biased than alternative strategies.

AlleleHMM is applicable to any type of functional genomic data and to any diploid species with a high-quality phased reference genome. AlleleHMM can now be deployed to understand the interplay between chromatin environment, transcription, and mRNA across a wide variety of organisms, providing new insights into how DNA sequences influence biochemical processes in the nucleus.

## Acknowledgements

We thank members of the Danko laboratory for valuable discussions. Work in this publication was supported by an NHGRI (National Human Genome Research Institute) grant R01-HG009309 to CGD. The content is solely the responsibility of the authors and does not necessarily represent the official views of the US National Institutes of Health.

---

Supplementary Figure 1: AlleleHMM had better sensitivity and specificity compared with using independent binomial tests for each SNP assuming a low heterozygosity (10 SNPs/ gene).

(A) Scatterplots show the sensitivity of AlleleHMM (red) and independent binomial tests (blue) as a function of the average read count at each SNP (left), or the expected maternal fraction (right). Error bars represent the standard error of 1000 independent simulations.

(B) Scatterplot shows the specificity of AlleleHMM (red) and independent binomial tests (blue) as a function of the average read count at each SNP. Error bars represent the standard error of 1000 independent simulations.

Supplementary Figure 2: The size of AlleleHMM blocks increased as read depth decreased.

(A) Histograms show the fraction of AlleleHMM blocks having a block size indicated on the X axis. Data is shown for the full GM12878 GRO-seq dataset (187,896,441 reads; blue), or a mock dataset subsample to 20 million reads (yellow).

(B) Histograms show the counts of AlleleHMM blocks as a function the block size. Blue ones were calculated using total GRO-seq reads (187,896,441) from GM12878, yellow ones were calculated using a subsample dataset of 20 million reads.

Supplementary Figure 3: AlleleHMM identifies blocks with fewer reads supporting each SNP. Histograms show the fraction of SNPs as a function the read counts per allele-specific SNPs identified by AlleleHMM only (H, red, top), independent binomial tests (I, blue, bottom), and the intersect of both methods (B, purple, middle) using GRO-seq data from GM12878.

